# Protection from α-synuclein-induced dopaminergic neurodegeneration by overexpression of the mitochondrial import receptor TOM20 in the rat midbrain

**DOI:** 10.1101/2020.05.11.089078

**Authors:** Briana R. De Miranda, Emily M. Rocha, Sandra Castro, J. Timothy Greenamyre

## Abstract

Dopaminergic neurons of the substantia nigra are selectively vulnerable to mitochondrial dysfunction, which is hypothesized to be an early and fundamental pathogenic mechanism in Parkinson’s disease (PD). Mitochondrial function depends on the successful import of nuclear-encoded proteins, many of which are transported through the TOM20-TOM22 outer mitochondrial membrane import receptor machinery. Recent data suggests that post-translational modifications of *α*-synuclein promote its interaction with TOM20 at the outer mitochondrial membrane and thereby inhibit normal protein import, which leads to dysfunction and death of dopaminergic neurons. As such, preservation of mitochondrial import in the face of α-synuclein accumulation might be a strategy to prevent dopaminergic neurodegeneration, however, this is difficult to assess using current *in vivo* models of PD. To this end, we established an exogenous co-expression system, utilizing AAV2 vectors to overexpress human α-synuclein and TOM20, individually or together, in the adult Lewis rat substantia nigra in order to assess whether TOM20 overexpression attenuates α-synuclein-induced dopaminergic neurodegeneration. Twelve weeks after viral injection, we observed that AAV2-TOM20 expression was sufficient to prevent loss of nigral dopaminergic neurons caused by AAV2-αSyn overexpression. The observed TOM20-mediated dopaminergic neuron preservation appeared to be due, in part, to the rescued import of nuclear-encoded mitochondrial electron transport chain proteins that were inhibited by α-synuclein overexpression. In addition, TOM20 overexpression rescued the import of the chaperone protein GRP75/mtHSP70/mortalin, a stress-response protein involved in α-synuclein-induced injury. Collectively, these data indicate that TOM20 expression prevents α-synuclein-induced mitochondrial dysfunction, which is sufficient to rescue dopaminergic neurons in the adult rat brain.

## Introduction

Among the characteristic molecular pathologies of Parkinson’s disease (PD) is the accumulation of α-synuclein within nigrostriatal dopaminergic neurons. In parallel, these neurons exhibit mitochondrial dysfunction which, in turn, has downstream consequences, including oxidative stress, inflammatory activation, impaired protein trafficking and degradation, and disrupted cellular signaling. Recently, we described a mechanism by which α-synuclein directly interacts with the mitochondrial translocase of the outer membrane (TOM) receptor, TOM20, and reduces import of proteins which contain an N-terminal mitochondrial targeting signal (MTS)^1^. These data showed that oligomeric, and post-translationally modified α-synuclein (oxidized, or dopamine modified), but not monomeric or nitrated α-synuclein, bind to TOM20 and prevent its association with TOM22, a key step in formation of the TOM complex that is necessary for protein import^1^. Because mitochondria must import approximately 99% of the proteins they contain^2,3^, blockade of mitochondrial protein import by toxic species of α-synuclein may be an early and important contributing factor to dopaminergic neurodegeneration^4,5^.

In rodent studies, overexpression of monomeric wildtype α-synuclein within the substantia nigra, either through viral-mediated expression of SNCA or direct α-synuclein particle seeding, results in degeneration of dopaminergic neurons and their terminal projections to the striatum^6-8^. α-Synuclein accumulation and mitochondrial dysfunction are both implicated as mechanisms that contribute to dopaminergic neurodegeneration in PD^4,9-12^. Thus, interaction between α-synuclein and TOM20-mediated protein import may be central to dopaminergic neuron dysfunction and death. Conversely, blocking this mechanism of cellular dysfunction may provide a therapeutic strategy to rescue dopaminergic neurons at an early point in disease pathogenesis.

To this end, we overexpressed human TOM20 in the substantia nigra pars compacta (SNpc) using an adeno-associated (AAV)2-TOM20 vector, injected unilaterally in adult Lewis rats. AAV2-TOM20 was co-expressed with either AAV2-GFP (control) or AAV2-αSyn. Similar to previous reports, AAV2-αSyn treatment caused significant loss of dopaminergic neurons following 12 weeks of viral incubation, and surviving cells contained both soluble and insoluble α-synuclein aggregates that correspond to human Lewy pathology^13-15^. Co-expression of AAV2-TOM20 with AAV2-αSyn did not reduce α-synuclein expression or accumulation within cells of the SNpc, however, it did result in neuroprotection against dopaminergic cell death. In conjunction, AAV2-TOM20 expression rescued mitochondrial import of nuclear-encoded proteins in this *in vivo*, proof-of-principal study examining the interaction of α-synuclein and mitochondrial function within dopaminergic neurons.

## Materials and Methods

### Adeno-associated viral vectors

Viral vectors were utilized to overexpress GFP, human wild type α-synuclein, or human TOM20. Each vector was expressed in an adeno-associated virus (serotype AAV2). The AAV2-TOM20 vector was produced by the Gene Therapy Program of the University of Pennsylvania Vector Core Program (Philadelphia, PA). AAV2-αSyn and AAV2-GFP vectors were purchased from the University of North Carolina Vector Core, through a partnership with the Michael J. Fox Foundation. Vector information is listed in **Table 1**.

**Table 1.**
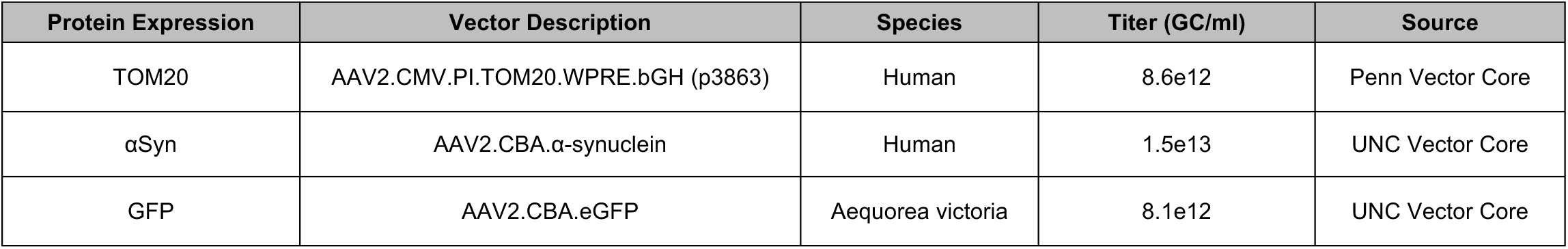
Vector information.

### Chemical reagents and supplies

Chemicals were purchased from Sigma-Aldrich (St. Louis, MO) unless otherwise noted. Antibodies are listed in **Table 2**.

**Table 2.** Antibody information.

| Antibody | Catalog | Tissue Concentration | Company |
| --- | --- | --- | --- |
| Tyrosine hydroxylase | AB1542 | 1:2000 | EMD Millipore (Burlington, MA) |
| Human-α-Synuclein (LB509) | Ab180215 | 1:1000 | Abcam (Cambridge, MA) |
| COX-IV | Ab16056 | 1:500 |  |
| GRP75/mtHSP70 | Ab2799 | 1:500 |  |
| TOM20 (Ms) | 612278 | 1:500 | BD Biosciences (San Jose, CA) |
| α-Synuclein (total) | 610787 | 1:500 | |
| NDUFS3/OxPhos | 459130 | 1:500 | Thermo Fisher (Waltham, MA) |
| TOM20 (Rb) | SC-11415 | 1:500 | Santa Cruz Biotechnology (Santa Cruz, CA) |
| SDHA | NBP1-71688 | 1:500 | Novus Biologicals (Centennial, CO) |

### Stereotaxic rodent surgery

Adult, middle age (9 month), male Lewis rats (Envigo) were utilized for AAV2 vector delivery. Animals were maintained under standard temperature and humidity-controlled conditions with 12:12 hour light-dark cycle. Conventional diet and water were provided *ad libitum*. Rats were randomly assigned to each treatment group and researchers were blinded to animal treatment throughout all behavioral analyses following AAV2 vector delivery.

Rodent stereotaxic surgery was performed under deep isoflurane anesthesia. Each animal received a unilateral infusion of two AAV2 vectors simultaneously; (1) GFP and TOM20, (2) GFP and αSyn, or (3) αSyn and TOM20, using standard Bregma coordinates for the substantia nigra (Bregma −5.8 mm A/P, −2.2 mm M/L, −8.5 mm V). Post-operative care included daily analgesic buprenorphine injections for three days following surgery, and sutures were removed upon wound closure. Animals were single-housed for the entirety of the 12-week study period, and euthanized using a lethal dose of pentobarbital, followed by transcardial perfusion with PBS and 4% paraformaldehyde perfusion fixation. All experiments involving animal treatment and euthanasia were approved by the University of Pittsburgh Institutional Animal Care and Use Committee.

### Motor behavior analysis

Prior to stereotactic surgery, animals were habituated to the postural instability test (PIT), and daily handling. PIT was used to assess asymmetric motor function, and is described in detail by Woodlee et al.^16^. Briefly, each animal was held vertically with one forelimb allowed to contact the table surface, which was lined with medium grit sandpaper. The animal’s center of gravity was then advanced until the animal triggered a “catch-up” step. The distance (cm) required for the animal to regain the center of gravity was recorded. Three trials were assessed per forelimb at each timepoint, and the average distance for each trial was recorded. PIT was assessed weekly for contralateral and ipsilateral forelimbs three weeks prior to surgery, and for 12-weeks post vector infusion. Behavioral tests were carried out by investigators blinded to treatment group throughout the study. Cylinder test recording was conducted at week 12; animals were placed in a glass cylinder (diameter 14 inches) and recorded for five continuous minutes in a closed environment. Video assessment of paw touches was carried out using blinded analysis.

### Rotenone administration

Adult (10 month) male and female Lewis rats (Envigo) were separated into single-housing, and handled for two weeks prior to the onset of rotenone administration. Rotenone was dissolved in DMSO (2% final concentration) and Miglyol 812 to reach the final concentration of 2.8 mg/kg. Rotenone handling and disposal was carried out following Unviersity of Pittsburgh Environmental Health and Safety procedures. Acute (5 Day ROT), endpoint rotenone (Endpoint ROT) and vehicle (2% DMSO and Miglyol) groups were randomly divided, and each animal was administered a single daily intraperitoneal (*i*.*p*.) injection of rotenone for 5 days (acute) or until they reached their motor behavioral endpoint, defined by the inability to perform the postural instability test or loss of 25% body mass.

### Striatal terminal intensity

Coronal rat brain sections (35 µm) encompassing the volume of the rat striatum (1/6 sampling fraction, approximately 10 sections per animal) were stained for tyrosine hydroxylase (TH) and detected using an infrared secondary antibody (IRDye® 680, LiCor Biosciences). Striatal tissue sections were analyzed using near-infrared imaging for density of dopamine neuron terminals (LiCor Odyssey) and analyzed using LiCor Odyssey software (V3.0; Licor Biosciences, Lincoln, NE). Output for striatal TH intensity is reported as arbitrary fluorescence units.

### Stereology

Stereological analysis of dopamine neuron number in the SN was achieved using an adapted protocol from Tapias et al. (2013)^17^ as reported in De Miranda et al. (2018, 2019)^18,19^ employing an unbiased, automated system. Briefly, nigral tissue sections were stained for TH and counterstained with DAPI and NeuroTrace Dye (640; Life Technologies) and imaged using a Nikon 90i upright fluorescence microscope equipped with high N.A. plan fluor/apochromat objectives, Renishaw linear encoded microscope stage (Prior Electronics) and Q-imaging Retiga cooled CCD camera (Center for Biological Imaging, University of Pittsburgh). Images were processed using Nikon NIS-Elements Advanced Research software (Version 4.5, Nikon, Melville, NY), and quantitative analysis was performed on fluorescent images colocalizing DAPI, TH, and Nissl-positive stains. Results are reported as the number of TH-positive cell bodies (whole neurons) within the SN.

### Immunohistochemistry and pathology

Brain sections (35 µm) were maintained at −20°C in cryoprotectant, stained while free-floating, and mounted to glass slides for imaging, using a “primary antibody delete” (secondary antibody only) stained section to establish background fluorescence limits. Fluorescent immunohistochemical images were collected using an Olympus BX61 confocal microscope and Fluoview 1000 software (Melville, NY). Quantitative fluorescence measurements were thoroughly monitored using standard operating imaging parameters to ensure that images contained no saturated pixels. For quantitative comparisons, all imaging parameters (e.g., laser power, exposure, pinhole) were held constant across specimens. Confocal images were analyzed using Nikon NIS-Elements Advanced Research software (Version 4.5, Nikon, Melville, NY). Mitochondrial colocalization of imported proteins (e.g. NDUFS3, SDHA, COX-IV) was determined by the boundary of TOM20 within transfected dopaminergic neurons, measured using Nikon NIS-Elements automated colocalization thresholding, to provide objective analysis. A minimum of 6 images per tissue slice were analyzed per animal, averaging 9-15 neurons per 60-100x image (approximately 180 cells per animal, per histological stain). 20x magnification was used to generate montage imaging of the ventral midbrain, for which the entire SN was analyzed per image using anatomical region of interest (ROI) boundaries. Results are reported as a measure of puncta within TH-positive cells, either number of objects (# of objects), or area per neuron in square pixels (px^2^), generated by Nikon NIS-Elements Advanced Research software.

### Statistical analyses

*A priori* power analysis was conducted to determine the minimal number of animals required to achieve 20-40% variance of the mean, with a 95% power at α=0.05 using G*Power statistical software (N=5 per group). All data were expressed as mean values ± standard error of the mean (SEM). Statistical significance was evaluated between normally distributed means by parametric one-way analysis of variance (ANOVA) with the Tukey *post-hoc* test to compare multiple data sets, or an unpaired t-test comparison of two means. Statistical significance is represented in each Figure as \**p* < 0.05, \*\**p* < 0.01, \*\*\**p* < 0.001, \*\*\*\**p* < 0.0001, unless otherwise specified on graph. F and T statistics are reported in each figure legend for ANOVA and t-tests, respectively. Postural instability was evaluated using the Kruskal-Wallis test (*p* < 0.05). Statistical analyses were carried out using GraphPad Prism software (V. 8.3.0).

## Results

### Characterization of AAV2-mediated TOM20 and α-synuclein overexpression

Adult (10-month-old) male Lewis rats were randomly assigned into three treatment groups and received a combination of two AAV2 viral vectors: AAV2-TOM20/GFP (control), AAV2-αSyn/GFP (disease control), or AAV2-TOM20/αSyn (**Figure 1A**). Following vector infusion, animals were monitored over a 12-week period, previously established for maximum AAV2-mediated α-synuclein expression to occur (**Figure 1B)**^20^. Midbrain histological analyses revealed a robust protein overexpression with each viral vector (human TOM20, GFP, or human αSyn) in the substantia nigra ipsilateral to the stereotactic injection (**Figure 1C-D**).

**Fig 1.**
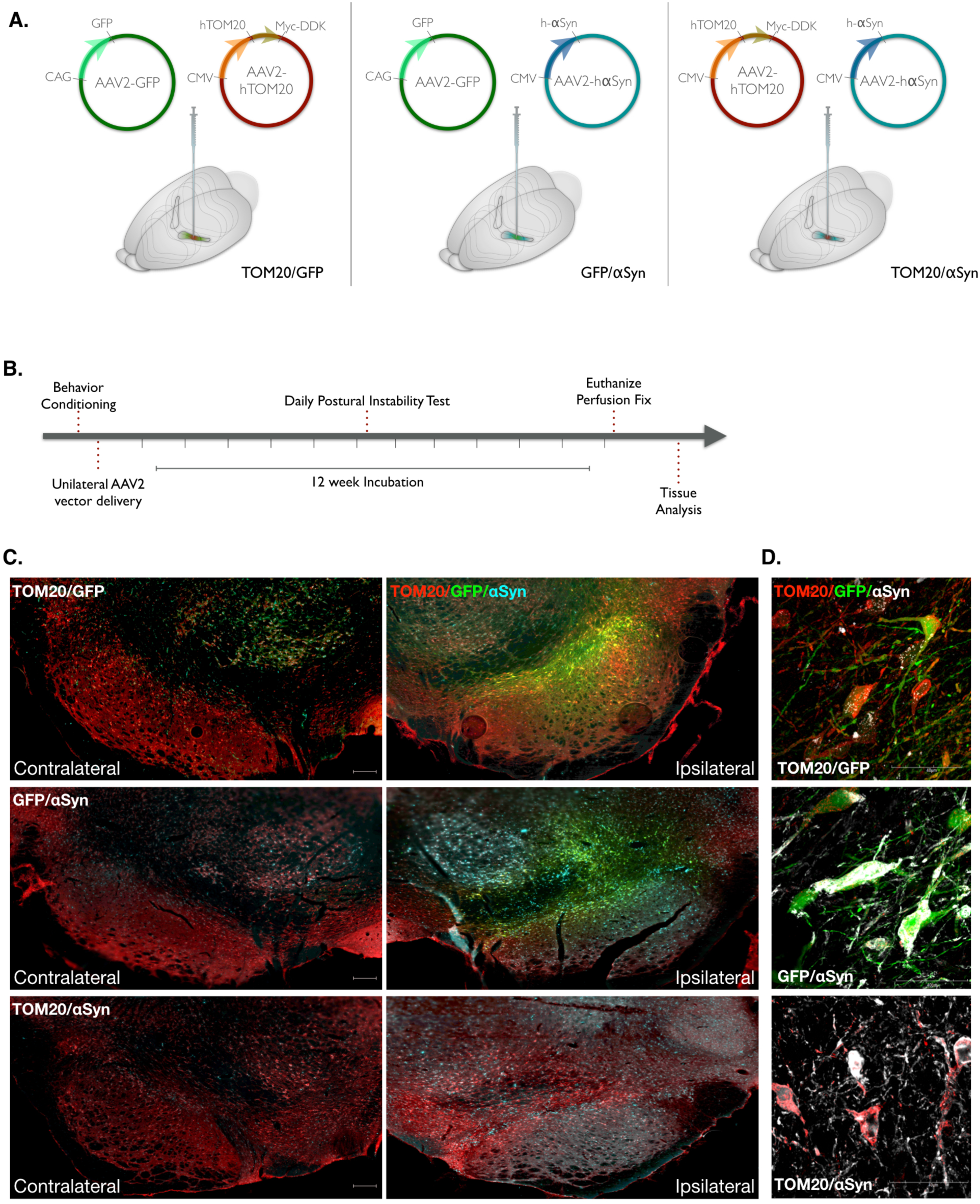
AAV2-mediated expression of human TOM20 and αSyn in the rat midbrain. AAV2 vectors driving protein overexpression were injected via stereotaxic surgery into the right substantia nigra of adult, male Lewis rats. **A**. Vector combinations of AAV2-TOM20/GFP, AAV2-αSyn/GFP, and AAV2-TOM20/αSyn represent the three treatment groups utilized within this study. **B**. Following vector infusion, animals were monitored daily for motor behavior, and euthanized for tissue collection 12-weeks post-injection. **C**. Representative images from each vector treatment group comparing ipsilateral (injected) and contralateral (non-injected) brain hemispheres; TOM20 (red), αSyn (cyan), GFP (green); scale bar 100μm. **D**. Target protein overexpression in neurons 12 weeks following vector expression; TOM20 (red), αSyn (white), GFP (green).

Animals that received AAV2-αSyn/GFP or AAV2-TOM20/αSyn injections displayed a robust expression of human α-synuclein protein with the neurons of the substantia nigra, which was absent in the AAV2-TOM20/GFP control group (**Figure 2A-B**). Midbrain tissue from animals expressing human AAV2-TOM20/αSyn was incubated with proteinase K, which revealed a fraction of insoluble α-synuclein protein aggregates following 12 weeks of viral vector incubation within the ipsilateral injection hemisphere (**Figure 2C**). Animals receiving AAV2-TOM20/GFP or AAV2-TOM20/αSyn vectors expressed approximately 5-fold increase of TOM20 protein within dopaminergic neuron cell bodies (positive for tyrosine hydroxylase; TH) compared to the noninjected (contralateral) hemisphere (**Figure 2D-E**).

**Fig 2.**
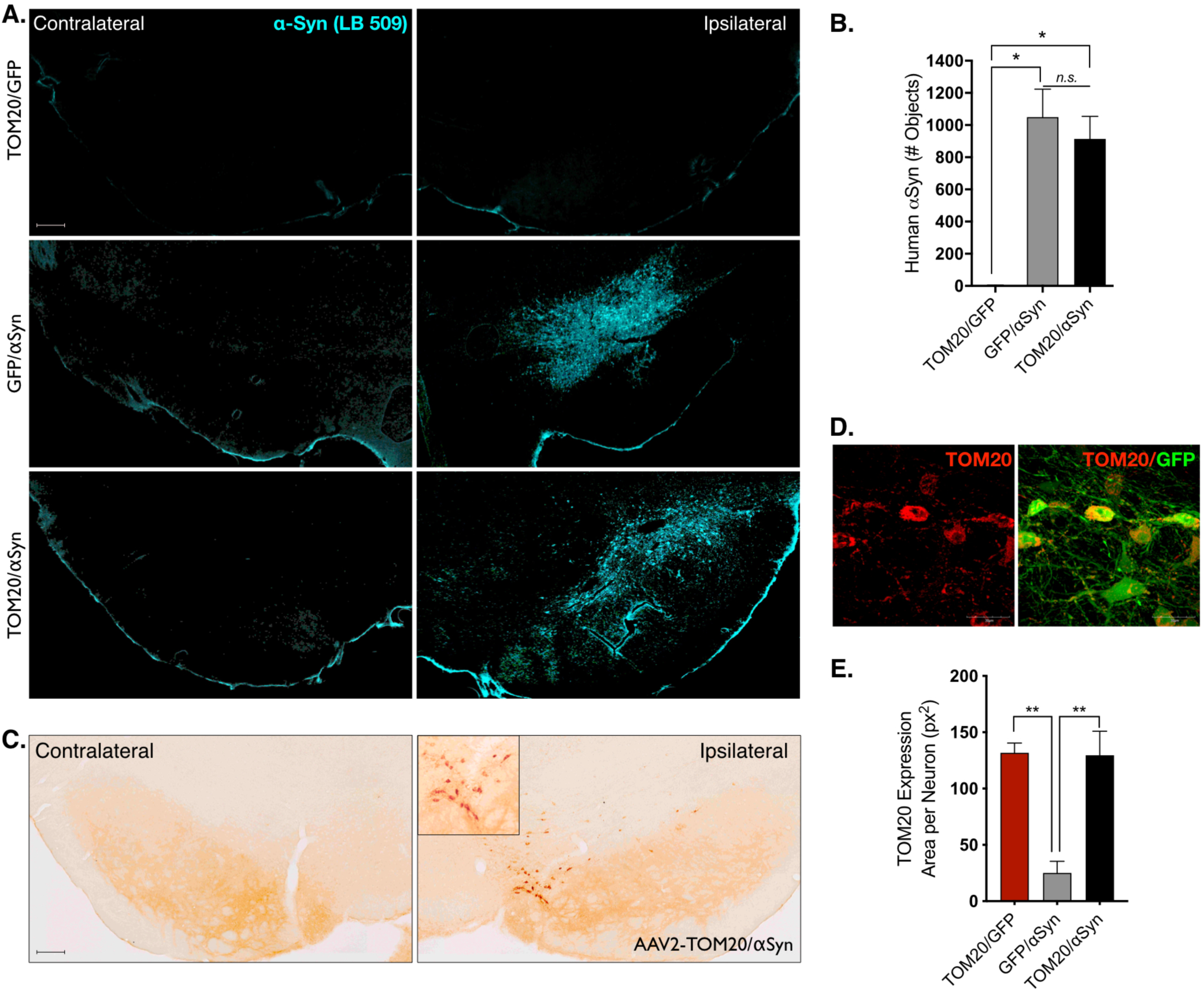
Vector co-administration does not substantially alter target protein expression. **A-B**. Human αSyn protein expression was detectable within the injected hemisphere of AAV2-αSyn/GFP and AAV2-TOM20/αSyn and was expressed at similar levels in both treatment groups receiving the human αSyn vector; (F(2, 26 = 4.6), *p* = 0.0192, ANOVA); scale bar 100μm. **C**. Insoluble αSyn (proteinase K resistant) was confirmed in animals injected with AAV2-αSyn vectors; representative image from AAV2-TOM20/αSyn; scale bar 100μm. **D-E**. AAV2-TOM20 vector infusion resulted in significantly elevated levels of TOM20 within neurons of the ventral midbrain, which was not affected by co-expression of AAV2-αSyn; (F(2, 6 = 17.5), *p* = 0.0031, ANOVA).

Vector-infused animals were assayed for motor behavioral changes during the 12-week incubation period using a postural instability test to identify unilateral motor movement deficits. Postural instability testing (PIT) revealed no significant difference between treatment groups over the course of the study (**Supplemental Figure 1A**). A cylinder test was performed at the end of the 12-week incubation period; however, neither the total number of rears nor preferential paw placement was significantly affected by α-synuclein or TOM20 overexpression (**Supplemental Figure 1B**). Similarly, no significant morbidity was observed in any treatment group during the twelve-week time course of the study.

### TOM20 overexpression is protective against α-synuclein-induced neurodegeneration

Striatal brain sections from injected animals were assessed for TH-positive terminal density and compared to the contralateral uninjected hemisphere. Animals receiving AAV2-αSyn/GFP vectors had ∼50% reduced TH fiber density ipsilateral to virus injection (*p* = 0.015), and this was prevented in animals that received TOM20 overexpression vector in conjunction with α-synuclein (AAV2-TOM20/αSyn; **Figure 3A-B**). Similarly, dopaminergic neurons within the SNpc were significantly depleted (47% loss) by AAV2-mediated overexpression of α-synuclein (AAV2-αSyn/GFP; **Figure 3C-D**); however, co-expression with TOM20 protected against α-synuclein-induced dopaminergic neuron cell death (*p* = 0.0001; AAV2-TOM20/αSyn). Thus, at the level of both the terminals and the cell bodies, TOM20 overexpression provided protection against *α*-synuclein.

**Fig 3.**
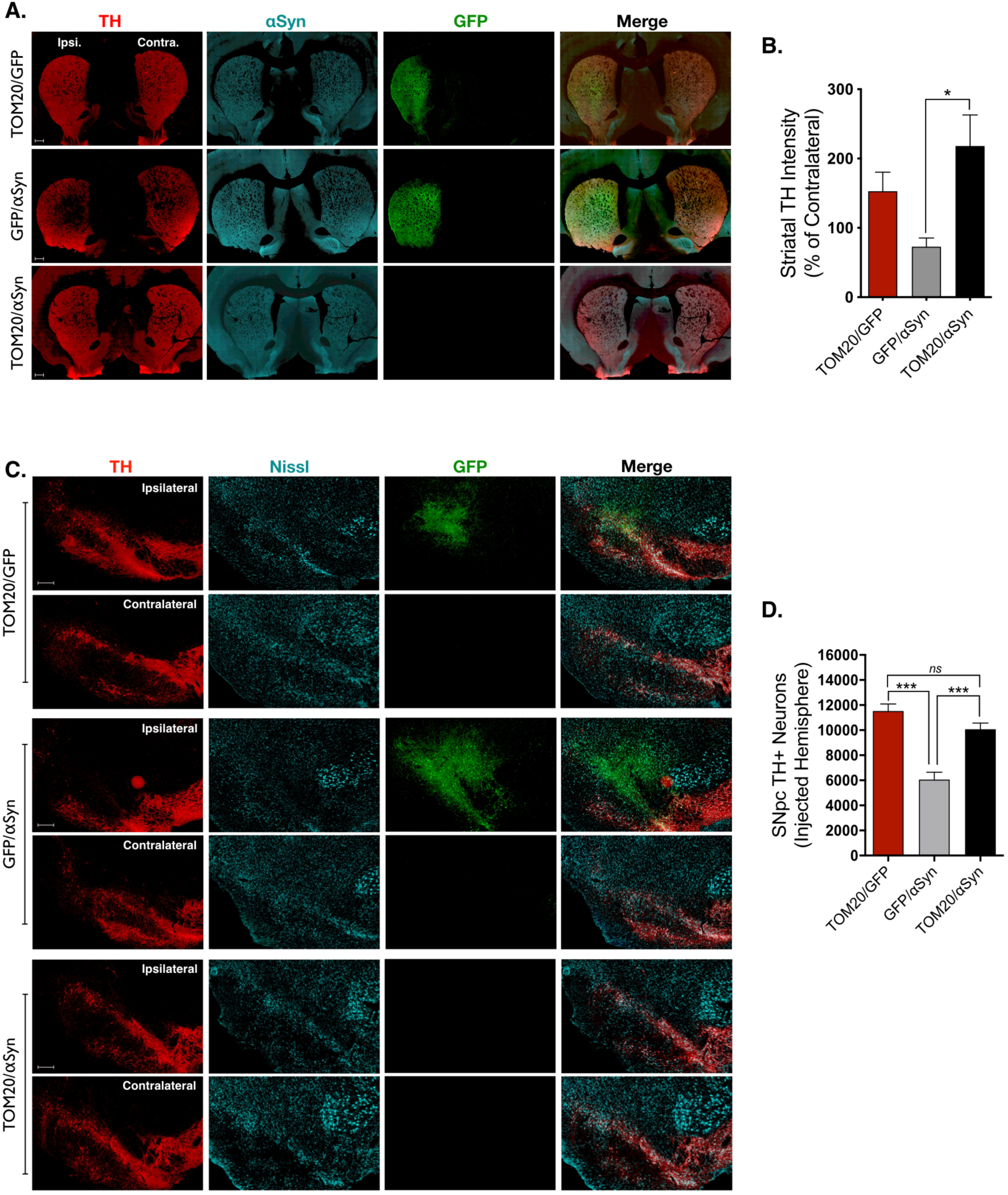
Overexpression of TOM20 was neuroprotective against AAV2-αSyn-mediated dopaminergic neurodegeneration. **A-B**. Dopaminergic terminals within the striatum (tyrosine hydroxylase; TH; red) were significantly reduced compared to the contralateral side in animals expressing AAV2-αSyn/GFP, but not in animals expressing AAV2-TOM20/αSyn; TH (red), αSyn (cyan), GFP (green); (F(2, 28 = 4.9), *p* = 0.0149, ANOVA). **C-D**. Representative images from the ipsilateral and contralateral hemispheres of each treatment group show significant dopaminergic neuron loss (TH, red) following AAV2-αSyn/GFP treatment, which was reduced by co-expression with TOM20 (AAV2-TOM20/αSyn); (F(2, 10 = 25.88), *p* = 0.0001, ANOVA); scale bars 100μm.

### Mitochondrial protein import impaired by α-synuclein is preserved by TOM20 overexpression

To determine whether TOM20 overexpression resulted in functional restoration of mitochondrial protein import, we assayed for nuclear encoded electron transport chain (ETC) proteins imported through the TOM20-TOM22 complex: NADH:Ubiquinone Oxidoreductase Core Subunit S3 (NDUFS3), succinate dehydrogenase [ubiquinone] flavoprotein subunit, mitochondrial (SDHA), and cytochrome c oxidase/complex IV (COX-IV). Following vector infusion, dopaminergic neurons in the SN that were transduced with AAV2-αSyn/GFP expressed significantly reduced protein levels of NDUFS3 (86% loss; *p* < 0.01), SDHA (90% loss; *p* < 0.0001), and COX-IV (85% loss; *p* < 0.05), compared to transduced neurons in AAV2-TOM20/GFP-injected animals (**Figure 4A-F**). In contrast, dopaminergic neurons that overexpressed both α-synuclein and TOM20 (AAV2-TOM20/αSyn) showed preserved mitochondrial import of NDUFS3 (p < 0.05), SHDA (*p* < 0.01), and COX-IV (*p* < 0.05).

**Fig 4.**
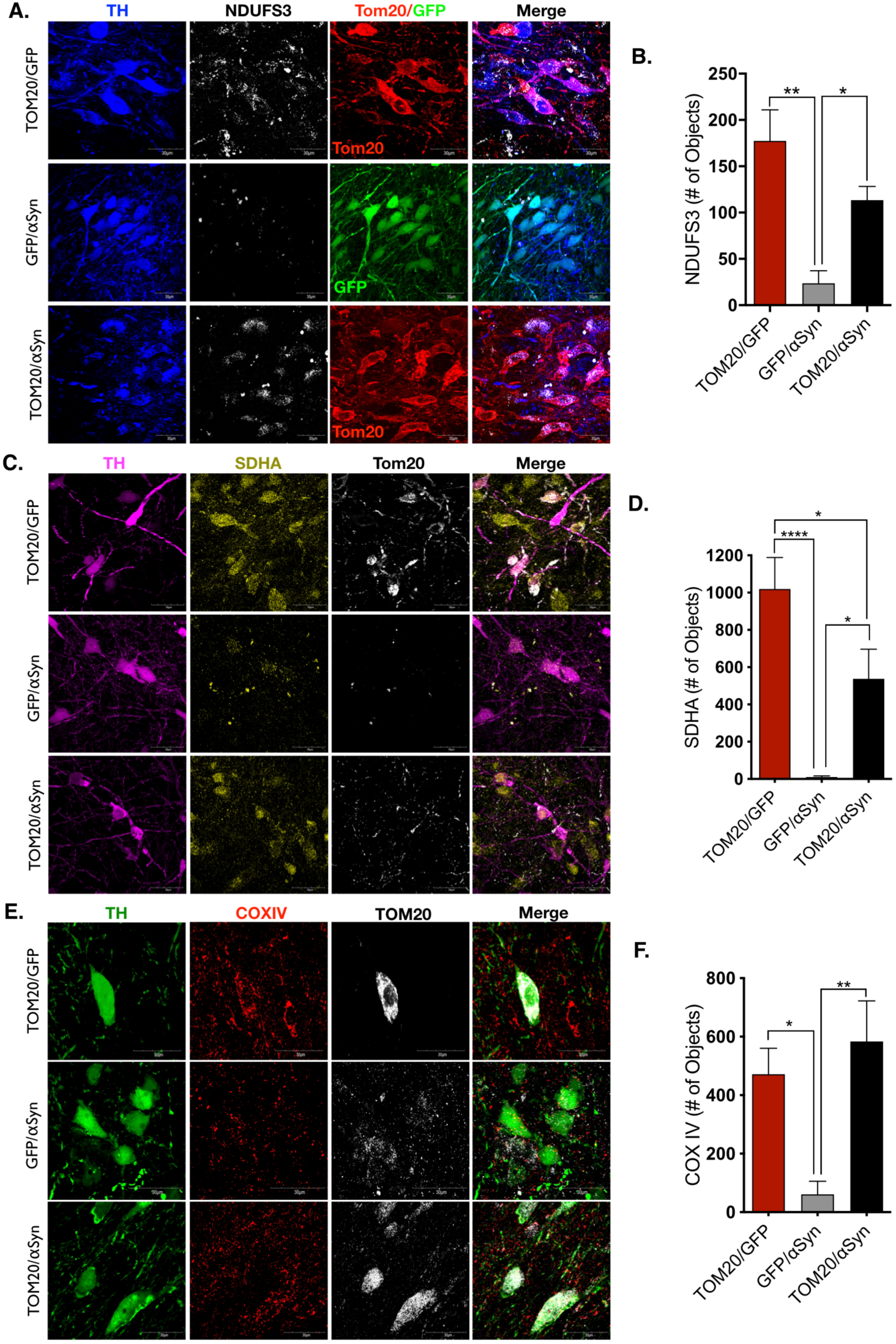
TOM20 expression rescued mitochondrial protein import impaired by αSyn. **A-B**. NADH:Ubiquinone Oxidoreductase Core Subunit S3 (NDUFS3, white) is a component of complex I of the electron transport chain. NDUFS3 import is significantly reduced in dopaminergic neurons of animals expressing AAV2-αSyn/GFP, and rescued by AAV2-TOM20 co-expression; TOM20 (red), GFP (green), TH (blue); (F(2, 10 = 11.82), *p* = 0.0023, ANOVA). **C-D**. Succinate dehydrogenase complex flavoprotein subunit A (SDHA, yellow) encodes the catalytic subunit of succinate-ubiquinone oxidoreductase of the electron transport chain, was reduced within mitochondria of dopaminergic neurons following αSyn overexpression, and rescued by AAV2-TOM20 co-expression; TOM20 (white), TH (magenta); (F(2, 23 = 14.57), *p* = 0.0001, ANOVA). **E-F**. Cytochrome c oxidase or complex IV (COX IV, red) the final enzyme in the mitochondrial transport chain, displayed a significant reduction within mitochondria in dopaminergic neurons expressing AAV2-αSyn/GFP, but not neurons expressing AAV2-TOM20/αSyn; TOM20 (red), TH (green); (F(2, 23 = 8.46), *p* = 0.0018, ANOVA).

### Mitochondrial import of GRP75/mtHSP70 expression is stress-responsive

Another nuclear-encoded mitochondrial protein, GRP75/mtHSP70 (mortalin), is both imported through the TOM20-TOM22 channel and is reportedly upregulated in response to mitochondrial oxidative stress, which can be caused by α-synuclein toxicity^4,21^. Unlike the imported ETC proteins, mitochondrial expression GRP75 was not significantly affected by α-synuclein overexpression (30% loss; *p* = 0.5), however, the combined expression of α-synuclein and TOM20 (AAV2-TOM20/αSyn) caused a robust increase in GRP75 (5-fold increase; *p* < 0.0001) that colocalized with mitochondria in dopaminergic neurons (**Figure 5A-B**). These data suggest that GRP75 imparts an important protective function in mitochondria damaged by α-synuclein accumulation, but GRP75 localization within the mitochondria is dependent on functional TOM20 import machinery.

**Fig 5.**
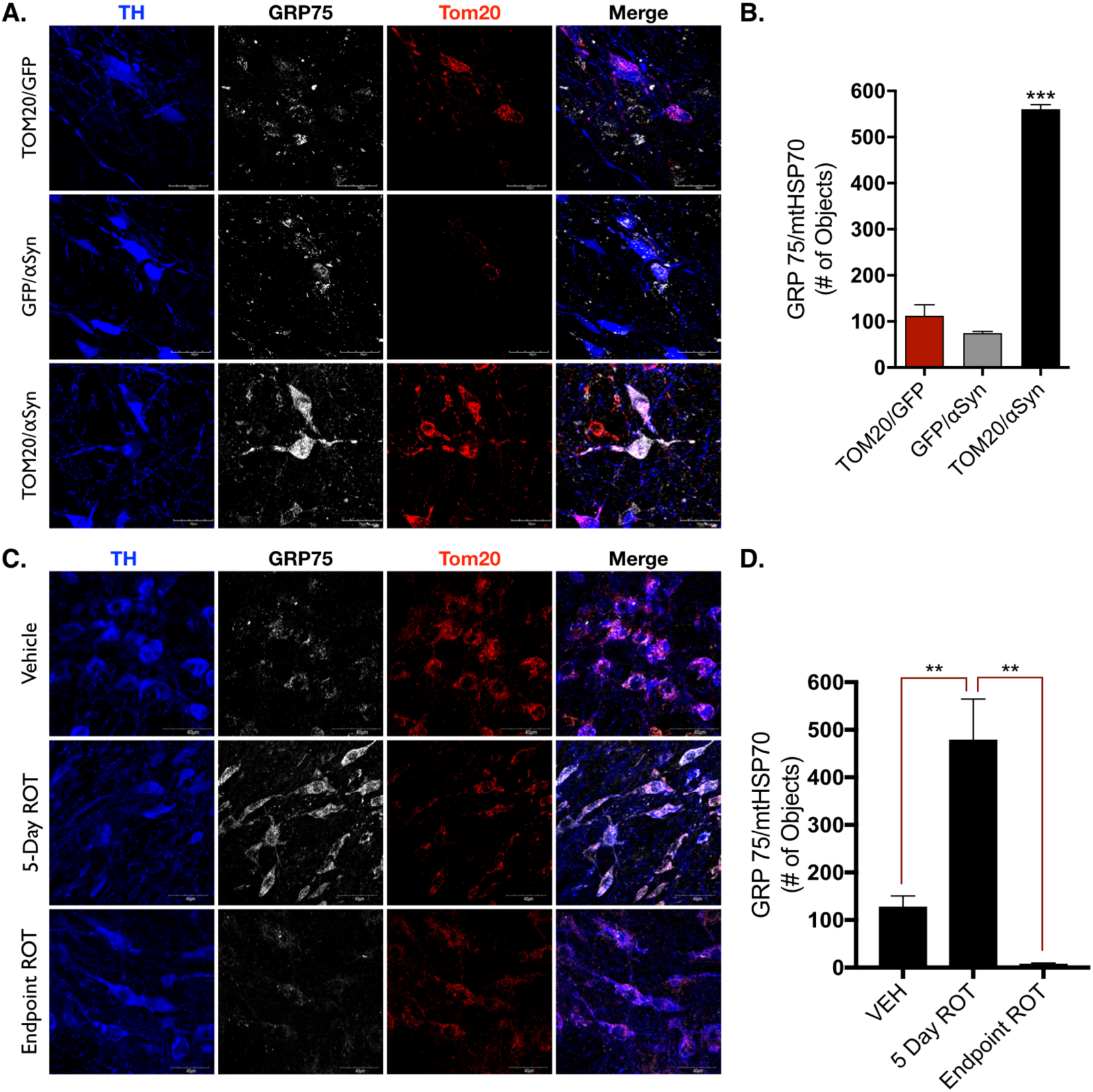
GRP75 expression is influenced by TOM20-mediated protein import. **A-B**. The redox-sensitive mitochondrial chaperone protein GRP75 was significantly elevated in dopaminergic neurons expressing AAV2-TOM20/αSyn, but not AAV2-αSyn/GFP, or AAV2-TOM20/GFP; (F(2, 4 = 167.2), *p* = 0.0001, ANOVA). **C-D**. Acute rotenone treatment causes the endogenous accumulation of αSyn (5-day ROT) that corresponded to a significant elevation in GRP75 expression within mitochondria, which was diminished in chronic/endpoint rotenone treated rats; (F(2, 15 = 14.69) *p* = 0.0003, ANOVA).

To further investigate the stress-dependent response of GRP75 import into the mitochondria, we treated adult (10 month) male Lewis rats with neutoroxin and prototypical complex I inhibitor rotenone, which causes significant mitochondrial dysfunction within dopaminergic neurons^22,23^. Animals were given a daily intraperitoneal (i.p.) rotenone injection (2.8 mg/kg) over an acute time course (5 days), which results in a marked oxidative stress response, and the accumulation of endogenous α-synuclein within dopaminergic neurons, but does not cause dopaminergic neurodegeneration^15^. A separate cohort of adult male Lewis rats were administered “endpoint” or chronic rotenone treatment, in which animals are given a daily injection of rotenone (2.8 mg/kg) until they exhibit predefined motor-behavioral impairment, which coincides with dopaminergic neuron death in the substantia nigra^19,24,25^. Acute (5 Day ROT) treatment resulted in a significant *increase* in GRP75 protein within dopaminergic neurons of the SN (*p* = 0.0011), but GRP75 expression was significantly *decreased* (*p* = 0.001) in dopaminergic neurons of animals receiving chronic rotenone treatment (Endpoint ROT; **Figure 5C-D**). Thus, elevated expression of GRP75 appears to be an early, possibly compensatory, response to mitochondrial stress, which cannot be sustained with prolonged or ongoing impairment, likely because of dysfunctional import into mitochondria.

## Discussion

Post-translational modification (PTM) and oligomerization of α-synuclein are implicated in the mitochondrial dysfunction that may precede dopaminergic neurodegeneration in PD^26^. These “toxic” forms of the protein have been observed in postmortem brain tissue of individuals with idiopathic PD using assays for oligomeric^27^ and phosphorylated forms of α-synuclein^12,28^. In this context, we previously reported that oligomeric or PTM forms of α-synuclein bind to the TOM20 subunit of the TOM20-TOM22 import complex and block mitochondrial protein import, a process which we proposed could be a possible target for therapeutic intervention to prevent dopaminergic neurodegeneration in PD^1^. To this end, we have utilized a viral vector-mediated model of human α-synuclein overexpression in the SN of adult Lewis rats to reproduce accumulation of α-synuclein protein within dopaminergic neurons. By co-expressing TOM20 together with *α*-synuclein, we attempted to ameliorate impaired mitochondrial protein import deficits and thereby rescue dopaminergic neurons from α-synuclein-induced neurodegeneration. Our results confirm that α-synuclein overexpression causes nigrostriatal dopaminergic neurodegeneration, which is associated with loss of nuclear-encoded mitochondrial proteins^1^. Here, we report the novel *in vivo* finding that TOM20 overexpression restores levels of nuclear-encoded, imported mitochondrial proteins in dopaminergic neurons, even in the face of continued *α*-synuclein overexpression, and protects these neurons against degeneration.

The mitochondrial genome encodes only 13 of the roughly 1,300 proteins contained in the organelle. Therefore, mitochondria depend on specific import mechanisms to acquire 99% of their protein constituents^29^. Levels of inner membrane/matrix respiratory chain components essential for proper mitochondrial function, such as the iron-sulfur subunit of complex I, NDUFS3, the complex IV subunit, COXIV, and the complex II flavoprotein subunit, SDHA, were all markedly reduced by α-synuclein overexpression – and this was associated with nigrostriatal degeneration. In contrast, the combined expression of exogenous TOM20 with *α*-synuclein largely prevented the loss of imported proteins and protected against *α*-synuclein toxicity. The survival of SN neurons co-expressing TOM20 with α-synuclein indicates that preservation of mitochondrial protein import was sufficient to protect against α-synuclein-induced dopaminergic neuron death.

Nuclear-encoded proteins destined for mitochondrial import via the TOM20-TOM22 system contain an MTS, also known as a presequence^30^. Such proteins are guided through the inner membrane to the matrix by the translocase of the inner membrane import channel in concert with the matrix chaperone, mtHSP70 (GRP75/mortalin), to maintain proteins in an unfolded state as they are processed by the mitochondrial processing peptidase (MPP)^31^. The function of GRP75 is pleiotropic; it is involved in protein import, but also plays a role in mitochondria-endoplasmic reticulum (ER) coupling^32^, where it appears to regulate Ca^2+^ transfer through the mitochondria-associated membrane (MAM)^33^. GRP75 has been implicated in the pathogenesis of PD, and was shown to be decreased in the postmortem brain tissue, and serum of individuals with PD compared to age-matched controls^34,35^. The robust GRP75 signal seen in dopaminergic neurons from animals co-expressing exogenous *α*-synuclein and TOM20 – but neither ontrol nor α-synuclein-overexpressing animals – indicates that (i) GRP75 is upregulated in response to α-synuclein induced stress, and (ii) expression of this chaperone protein also requires functional mitochondrial import. This is further corroborated by our finding of elevated levels of GRP75 with acute rotenone, but dramatic loss of GRP75 with chronic rotenone exposure, at a time when mitochondrial import is known to be impaired^1^.

There were limitations of this study. The experimental design focused on histological outcomes; as such, the use of fixed tissue precluded assays of mitochondrial function and dynamics within dopaminergic neurons. Additionally, we found no behavioral deficits associated with *α*-synuclein overexpression and could therefore not measure improvement with TOM20 overexpression. This lack of a gross motor phenotype was most likely a result of the combination of a slowly evolving lesion (with ongoing compensation) and the moderate final level of nigrostriatal degeneration (∼50%).

Collectively, however, these data show that *α*-synuclein overexpression in dopaminergic neurons of the adult SN impairs mitochondrial protein import and causes neurodegeneration. TOM20 overexpression was sufficient to rescue α-synuclein-induced dopaminergic neurodegeneration over this 12-week time period, likely because it rescued mitochondrial function^1^, which is integral to dopaminergic neuron survival^36-38^. Additionally, these data suggest a role for the mitochondrial chaperone and redox sensitive protein, GRP75/mtHSP70/mortalin, in protection of dopaminergic neurons from α-synuclein induced pathology. Although multiple mechanisms of toxicity have been ascribed to α-synuclein, our finding of virtually complete neuroprotection with TOM20 overexpression indicates that its interactions with mitochondria may be a major means by which it causes neurodegeneration. As such, the α-synuclein-TOM20 interaction may represent an important target for therapeutic intervention.

## Acknowledgments

This work was supported by research grants from the National Institutes of Health (NS100744, R21ES027470, NS095387, and K99ES029986), the American Parkinson Disease Association, the Parkinson’s Foundation, the Michael J Fox Foundation, the Blechman Foundation, and the friends and family of Sean Logan.

---

**Supplemental Fig 1.**
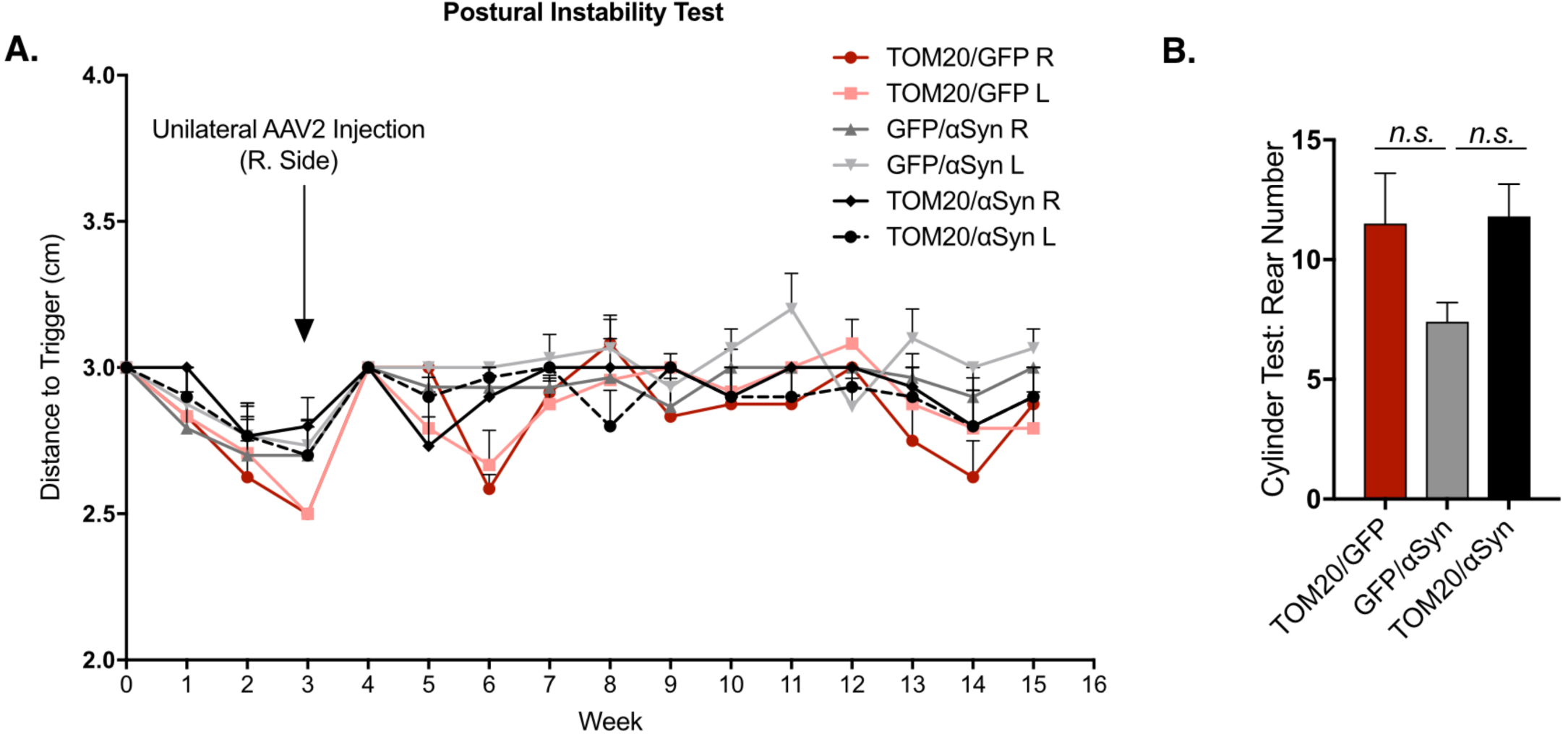
AAV2-αSyn expression within the substantia nigra did not result in significant behavioral abnormalities. **A**. The postural instability test (PIT) was performed at baseline (prior to vector infusion) and over the 12-week study timecourse. **B**. The average number of rearing movements from the cylinder test, performed at week 12; (F(2, 11 = 2.5) *p* = 0.1266, ANOVA.)

## References

1. Di Maio, R. et al. α-Synuclein binds to TOM20 and inhibits mitochondrial protein import in Parkinson’s disease. Science Translational Medicine 8, 342ra78–342ra78 (2016).

2. Anderson, S. et al. Sequence and organization of the human mitochondrial genome. Nature 290, 457–465 (1981).

3. Andrews, R. M. et al. Reanalysis and revision of the Cambridge reference sequence for human mitochondrial DNA. Nat Genet 23, 147–147 (1999).

4. Devi, L., Raghavendran, V., Prabhu, B. M., Avadhani, N. G. & Anandatheerthavarada, H. K. Mitochondrial import and accumulation of alpha-synuclein impair complex I in human dopaminergic neuronal cultures and Parkinson disease brain. J. Biol. Chem. 283, 9089–9100 (2008).

5. Franco-Iborra, S. et al. Defective mitochondrial protein import contributes to complex I-induced mitochondrial dysfunction and neurodegeneration in Parkinson’s disease. Cell Death and Disease 9, 1122–17 (2018).

6. Thakur, P. et al. Modeling Parkinson’s disease pathology by combination of fibril seeds and α-synuclein overexpression in the rat brain. Proc. Natl. Acad. Sci. U.S.A. 114, E8284–E8293 (2017).

7. Rocha, E. M. et al. Glucocerebrosidase gene therapy prevents α-synucleinopathy of midbrain dopamine neurons. Neurobiology of Disease 82, 495–503 (2015).

8. Volpicelli-Daley, L. A. et al. Exogenous α-synuclein fibrils induce Lewy body pathology leading to synaptic dysfunction and neuron death. Neuron 72, 57–71 (2011).

9. Rocha, E. M., De Miranda, B. & Sanders, L. H. Alpha-synuclein: Pathology, mitochondrial dysfunction and neuroinflammation in Parkinson’s disease. Neurobiology of Disease (2017).

10. Hsu, L. J. et al. alpha-synuclein promotes mitochondrial deficit and oxidative stress. The American Journal of Pathology 157, 401–410 (2000).

11. Smith, W. W. et al. Endoplasmic reticulum stress and mitochondrial cell death pathways mediate A53T mutant alpha-synuclein-induced toxicity. Human Molecular Genetics 14, 3801–3811 (2005).

12. Grassi, D. et al. Identification of a highly neurotoxic α-synuclein species inducing mitochondrial damage and mitophagy in Parkinson’s disease. Proc. Natl. Acad. Sci. U.S.A. 115, E2634–E2643 (2018).

13. Theodore, S., Cao, S., McLean, P. J. & Standaert, D. G. Targeted overexpression of human alpha-synuclein triggers microglial activation and an adaptive immune response in a mouse model of Parkinson disease. J Neuropathol Exp Neurol 67, 1149–1158 (2008).

14. St Martin, J. L. et al. Dopaminergic neuron loss and up-regulation of chaperone protein mRNA induced by targeted over-expression of alpha-synuclein in mouse substantia nigra. Journal of Neurochemistry 100, 1449–1457 (2007).

15. Di Maio, R. et al. LRRK2 activation in idiopathic Parkinson’s disease. Science Translational Medicine 10, eaar5429 (2018).

16. Woodlee, M. T., Kane, J. R., Chang, J., Cormack, L. K. & Schallert, T. Enhanced function in the good forelimb of hemi-parkinson rats: compensatory adaptation for contralateral postural instability? Experimental Neurology 211, 511–517 (2008).

17. Tapias, V., Greenamyre, J. T. & Watkins, S. C. Automated imaging system for fast quantitation of neurons, cell morphology and neurite morphometry in vivo and in vitro. Neurobiology of Disease 54, 158–168 (2013).

18. De Miranda, B. R. et al. Astrocyte-specific DJ-1 overexpression protects against rotenone-induced neurotoxicity in a rat model of Parkinson’s disease. Neurobiology of Disease 115, 101–114 (2018).

19. De Miranda, B. R., Fazzari, M., Rocha, E. M., Castro, S. & Greenamyre, J. T. Sex differences in rotenone sensitivity reflect the male-to-female ratio in human Parkinson’s disease incidence. Toxicol. Sci. 354, 319 (2019).

20. Zharikov, A. D. et al. shRNA targeting α-synuclein prevents neurodegeneration in a Parkinson’s disease model. J. Clin. Invest. 125, 2721–2735 (2015).

21. Liu, F.-T. et al. Involvement of mortalin/GRP75/mthsp70 in the mitochondrial impairments induced by A53T mutant α-synuclein. Brain Research 1604, 52–61 (2015).

22. Higgins, D. S. & Greenamyre, J. T. [3H]dihydrorotenone binding to NADH: ubiquinone reductase (complex I) of the electron transport chain: an autoradiographic study. Journal of Neuroscience 16, 3807–3816 (1996).

23. Testa, C. M., Sherer, T. B. & Greenamyre, J. T. Rotenone induces oxidative stress and dopaminergic neuron damage in organotypic substantia nigra cultures. Molecular Brain Research 134, 109–118 (2005).

24. Cannon, J. R. et al. A highly reproducible rotenone model of Parkinson’s disease. Neurobiology of Disease 34, 279–290 (2009).

25. Rocha, E. M. et al. LRRK2 inhibition prevents endolysosomal deficits seen in human Parkinson’s disease. Neurobiology of Disease 134, 104626 (2019).

26. Barrett, P. J. & Timothy Greenamyre, J. Post-translational modification of α-synuclein in Parkinson’s disease. Brain Research 1628, 247–253 (2015).

27. Roberts, R. F., Bengoa-Vergniory, N. & Alegre-Abarrategui, J. Alpha-Synuclein Proximity Ligation Assay (AS-PLA) in Brain Sections to Probe for Alpha-Synuclein Oligomers. Methods Mol. Biol. 1948, 69–76 (2019).

28. Grassi, D., Diaz-Perez, N., Volpicelli-Daley, L. A. & Lasmézas, C. I. Pα-syn* mitotoxicity is linked to MAPK activation and involves tau phosphorylation and aggregation at the mitochondria. Neurobiology of Disease 124, 248–262 (2019).

29. Copeland, W. C. & Longley, M. J. Mitochondrial genome maintenance in health and disease. DNA Repair 19, 190–198 (2014).

30. Omura, T. Mitochondria-targeting sequence, a multi-role sorting sequence recognized at all steps of protein import into mitochondria. J. Biochem. 123, 1010–1016 (1998).

31. Wiedemann, N., Frazier, A. E. & Pfanner, N. The protein import machinery of mitochondria. J. Biol. Chem. 279, 14473–14476 (2004).

32. Honrath, B., Culmsee, C. & Dolga, A. M. One protein, different cell fate: the differential outcome of depleting GRP75 during oxidative stress in neurons. Cell Death and Disease 9, 32–3 (2018).

33. Honrath, B. et al. Glucose-regulated protein 75 determines ER-mitochondrial coupling and sensitivity to oxidative stress in neuronal cells. Cell Death Discov 3, 17076–13 (2017).

34. Jin, J. et al. Proteomic identification of a stress protein, mortalin/mthsp70/GRP75: relevance to Parkinson disease. Mol. Cell Proteomics 5, 1193–1204 (2006).

35. Singh, A. P. et al. Serum Mortalin Correlated with α-Synuclein as Serum Markers in Parkinson’s Disease: A Pilot Study. Neuromolecular Med. 20, 83–89 (2018).

36. Bose, A. & Beal, M. F. Mitochondrial dysfunction in Parkinson’s disease. Journal of Neurochemistry 139 Suppl 1, 216–231 (2016).

37. Greenamyre, J. T., MacKenzie, G., Peng, T.-I. & Stephans, S. E. Mitochondrial dysfunction in Parkinson’s disease. Biochemical Society Symposia 66, 85–97 (1999).

38. Schapira, A. H. et al. Mitochondrial complex I deficiency in Parkinson’s disease. The Lancet 1, 1269 (1989).

